# Soil sanctuaries: Experimental manipulations enhance ground-nesting bee habitat across an urban gradient

**DOI:** 10.1101/2025.04.18.649528

**Authors:** A. Fredenburg, K. Goodell

## Abstract

1. As evidence of pollinator declines mounts, effective conservation strategies are urgently needed. Current methods focus on planting flowers and frequently overlook nesting habitats, especially for ground nesting bees, which comprise most species. Despite their prevalence, ground-nesting bee habitats remain poorly understood. The lack of evidence-based methods to enhance nesting habitat presents a significant challenge, particularly in environments with limited natural nesting sites, such as urban landscapes.
2. To determine whether soil manipulations enhance bee colonization rates, we constructed experimental nesting plots at 21 sites along an urban gradient and implemented five soil surface treatments: bare ground, leaf litter, mounded soil, pebbles, and unmanipulated controls. We collected nesting individuals from experimental plots through observation and emergence traps to investigate preferences for soil surface substrates, and the impact of plot-level and site-level conditions on colonization.
3. Manipulated treatments significantly increased colonization compared to controls and revealed genus-specific preferences. Treatments also resulted in differences in environmental conditions including temperature, moisture, and soil compaction.
4. Site-level conditions including slope and urbanization increased colonization while hard soil compaction and bare ground decreased colonization success. Urban and highly vegetated sites were more frequently colonized by ground nesting bees, indicating that our manipulations successfully attracted nest-searching females and may have alleviated limited nesting opportunities.
5. *Policy Implications:* Simple soil surface manipulations, or soil sanctuaries, represent a practical tool to enhance an essential component of bee habitat and provide a safe nesting location that can improve conservation outcomes for ground-nesting bees across diverse environments. Incorporating ground-nesting bee habitat enhancements into urban planning and land management policies could address habitat limitations for many species. Conservation strategies should prioritize the inclusion of nesting features alongside floral resources to create comprehensive pollinator-supportive landscapes, benefiting both biodiversity and critical ecosystem services.

## 1 Introduction

Pollinators are essential to terrestrial ecosystems, but anthropogenic disturbance is causing population declines (Potts et al., 2010). Disturbed habitats, such as urban areas, offer only small, fragmented patches of suitable habitat, making pollinator populations vulnerable. However, key pollinator taxa like bees, include many small species that can persist in these patches if local conditions fulfill their habitat requirements (Burks & Philpott, 2017). Therefore, small high-quality habitat patches, even in disturbed landscapes, have the potential to meet species’ ecological needs and serve as important refuges if properly managed (Hall et al., 2017). Pollinator conservation requires evidence-based habitat enhancements, particularly in areas with limited resources.

Disturbed habitats are often planted with flowers to attract pollinators, but these floral patches may not fulfill species requirements (Egerer et al., 2019). Wild bees require floral resources to persist (Potts et al., 2003), but they also need nesting sites and materials (Roulston & Goodell, 2011). As central place foragers, bees return to their nests between foraging bouts (Westrich, 1996). Thus, populations will only respond to increased floral resources if suitable nesting sites are within their foraging range of a few hundred meters (Zurbuchen et al., 2010). Nest location is crucial for both access to floral resources and protecting developing brood from flooding, parasitism, and other threats (Antoine and Forrest, 2021).

Many bee species exhibit philopatry, often nesting near their natal nests (Yanega, 1990). Therefore, providing safe spaces that have suitable nesting conditions, protected from disturbances, can promote long-term population growth (Roulston & Goodell, 2011). In disturbed areas, suitable nesting sites are likely limited, but habitat enhancement efforts often overlook nesting habitats (Harmon-Threatt, 2020). Effective bee conservation must prioritize enhancing both floral and nesting habitats.

Significant gaps in our understanding of bee nesting ecology hinder efforts to enhance nesting habitat. For species that nest above ground, enhancements such as providing reed stems or small paper tubes have successfully increased nest site availability (Steffan-Dewenter & Schiele, 2008). However, 80% of bee species nest in the soil and their nesting requirements remain largely unknown. As a result, effective means of supplementing nesting habitat have not yet been established (Leonard & Harmon-Threatt, 2019). Some studies have tested soil manipulations by constructing patches of bare soil (Gregory & Wright, 2005; Fortel et al., 2016; Gardein et al., 2022; Tsiolis et al., 2022) and adding features like pebbles (Cane, 2015), successfully providing nest sites for some species. However, controlled experiments on ground-nesting bee ecology are uncommon, even though they are essential for evaluating enhancement strategies (Orr, et al., 2022). Evidence shows that ground-nesting genera exhibit different nesting preferences (Sardiñas & Kremen, 2014; Maher et al., 2019), indicating that no single approach will support all species. Therefore, assessing a range of soil manipulations is critical for ground-nesting bee conservation.

The effectiveness of nesting habitat augmentation likely depends on the local availability of nesting sites and floral abundance. In highly disturbed areas like urban landscapes with extensive impervious surfaces that prevent nest excavation (Sivakoff et al., 2018), nest site enhancements may be particularly beneficial. Because bare ground availability has been associated with increased species richness (Potts et al., 2005), there is an assumption that it supports all ground-nesting bees. However, bare ground is only one of several potentially attractive nesting substrates and may only benefit specific taxa, such as Halictids (Fortel et al., 2016). Other substrates such as rocks (Cane, 2015), leaf litter (Davis & LaBerge, 1975), and features like slopes (Maher et al., 2019) may be equally, or more suitable for different species. Various substrates may attract bees through suitable environmental soil conditions including temperature, moisture, and compaction, as well as provide visual cues and protection from cleptoparasites (Antoine and Forrest, 2021). Thus, if we can provide a variety of soil surface substrates, we might be able to support diverse bee communities. Currently, we lack consensus on evidence-based guidance for enhancing ground-nesting bee habitats, but experimental results will build on anecdotal and observational findings to develop data-driven protocols.

In this study, we experimentally manipulated soil surface features at sites across an urban gradient to test their attractiveness to ground-nesting bees. We hypothesized that (1) Bee colonization would be higher in manipulated treatments due to their influence on environmental conditions, with bee genera preferring different treatments and (2) bee colonization would vary with site-level local habitat conditions and landscape-scale urbanization. We predicted colonization would increase with urbanization due to limited nest site availability, and local factors including bare ground, floral availability, slope, soil texture, and compaction would influence colonization success. We examine how manipulating natural substrates can provide nesting sites for local bee populations and how plot and site-level conditions influence nesting activity, providing novel insights into habitat enhancement strategies for ground-nesting bees.

## 2 Materials and Methods

### 2.1 Site Selection

We selected 21 sites (Table S1) across an urbanization gradient (1.3% – 88.0%) using the National Land Cover Database (Dewitz, 2021). Urbanization was calculated as the proportion of developed land (low, medium and high intensity) within a 500-m buffer centered on the experimental plot using ArcGIS Pro (ESRI, 2023). Sites consisted of city and county parks or urban greenspaces located in the northcentral USA in Columbus, Ohio. All sites included a flower patch ≥ 25 m^2^ within 200 m of the plot to ensuring forage access for even small bee species (Gathmann & Tscharntke, 2002). Distance to the flower patch was included as a site-level covariate.

### 2.2 Plot Construction

At each site, we constructed a 3-m x 5-m plot divided into three 1-m x 5-m replicate strips, each containing five 1-m^2^ subplots assigned to one of five treatments: bare ground, leaf litter, pebbles, mounded soil, or an unmanipulated control (Fig. 1a & b).

**Figure 1.**
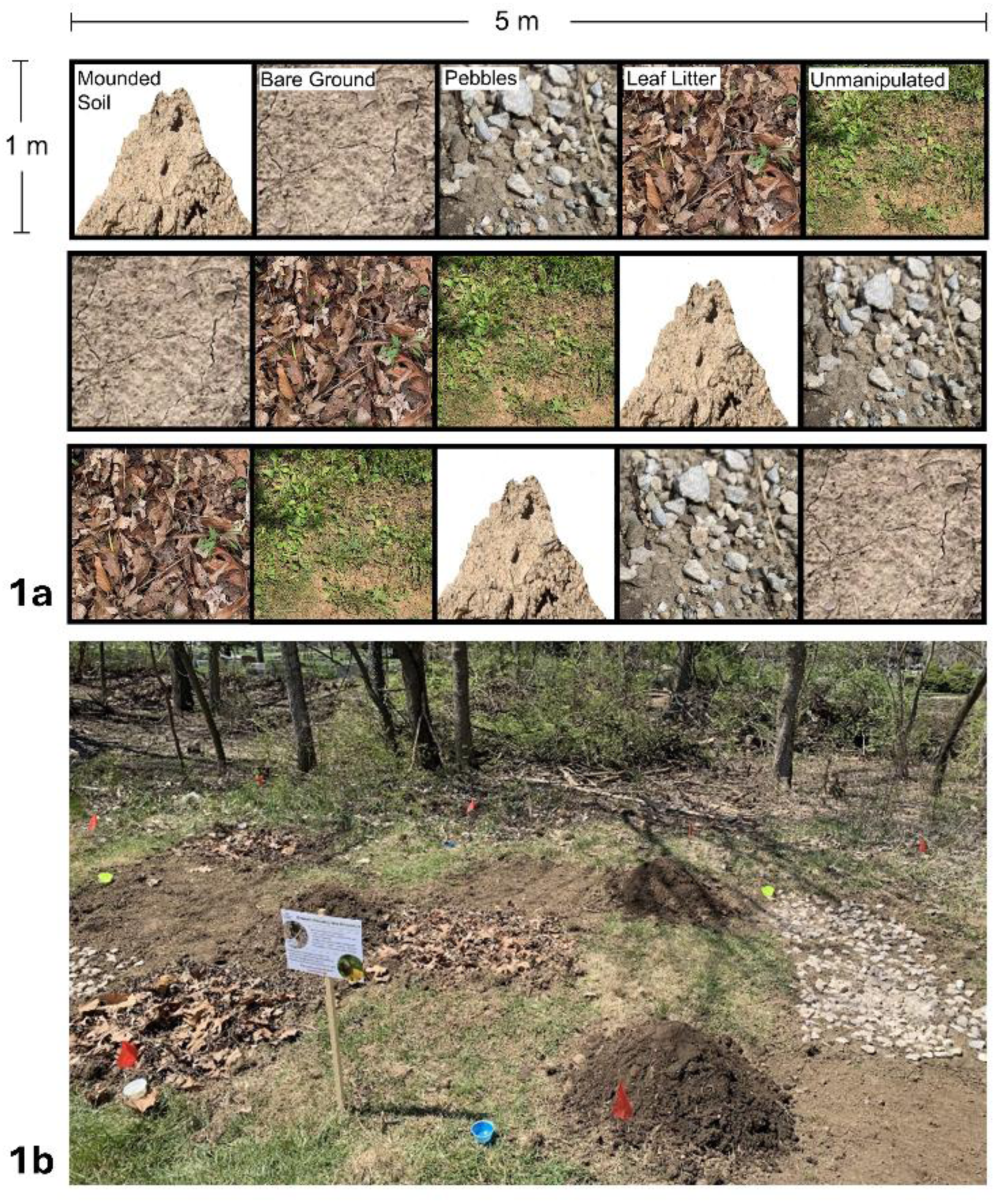
**a.** Visual representation of the five treatments arranged randomly in the three replicates at each site. **b.** photograph of a constructed experimental plot.

In March 2023, before bee emergence, we removed vegetation from manipulated treatments using a Honda Rear Tine Tiller (FRC800K1A) with 12″ (∼30 cm) diameter tines and standardized vegetation in control treatments to < 0.25 m high. In leaf litter treatments, we added a 2-cm layer of site-sourced dead leaves overlain with branches and in pebble treatments we covered 50% of the surface with drainage rocks (Cane, 2015). For mounded soil treatments, we constructed 25-cm-high soil mounds, with ∼2.0 kg/cm^2^ compacted soil (preferred compaction found by Kim et al., 2006), to simulate slopes in otherwise flat locations. We placed plots in open sunny areas on mostly flat ground and measured slope with iPhone application Bubble Level (Version 5.6) to use as a covariate. We placed 10 empty sauce cups painted blue, yellow, and white on the edges of the plot, drilled with drainage holes, and anchored with golf tees to attract bees and encourage habitat discovery.

### 2.3 Environmental Data Collection

We visited sites six times (∼once per month) from May–September 2023 and April 2024. During each visit, we recorded soil compaction and temperature from each of the 15 1-m x 1-m treatments. We used an infrared thermometer (Etekcity, Anaheim, USA) for temperature and a Pocket Penetrometer (Forestry Suppliers, Inc., Jackson, USA) for soil compaction. Because the penetrometer’s maximum reading was 4.5 kg/cm^2^, we categorized compaction as soft (< 1.9 kg/cm^2^), medium (2–3.9 kg/cm^2^), or hard (4 < kg/cm^2^) and counted the number of readings (15 total per visit) in each category, similarly done by Potts et al. (2005). We used a 36-in (91.4-cm) soil corer (JMC Soil Samplers, Newton, USA) to collect samples for lab-based soil texture analysis via laser diffraction (Faé et al. 2019). We also collected a ∼5 cm^2^ soil sample from the center of each treatment type in a randomly selected replicate using a trowel, refilling the hole to avoid affecting bee nesting. We measured soil wet weight, air dried for one week, and weighed again to calculate moisture content.

During each visit, we assessed site-level resources by measuring bare ground coverage, floral abundance, and distance to the nearest floral patch. We estimated % bare ground in five random quadrats along a 50-m transect extending from the plots in a random cardinal direction. For floral abundance, we counted the number of blooming floral units per plant species in five random quadrats along a 50-m transect centered on a flowering area within 200 m of the plot. We measured the distance to the nearest floral patch > 25 m^2^ from the plot edge. To estimate ground-nesting bees foraging at a site, we non-lethally netted bees for 15 minutes, tallied by genus and temporarily placed them in a jar inside a cooler to avoid recounts, then released them after the survey.

### 2.4 Nesting Bee Data Collection

We captured nesting bees using two methods. First, we observed the plot for 15 minutes and lethally collecting bees entering and exiting nesting, also collecting opportunistically collected bees throughout the ∼1.5-hour site visits. Second, we deployed emergence traps in May and July 2023 and April 2024. Traps, modified from Cadmus et al. (2016), consisted of a vertical 50*-*cm half-inch PVC pipe placed upright at the center of each 1-m^2^ subplot, covered with fine mesh, secured by a 1-m^2^ PVC frame anchored with landscape fabric pins, and held flush to the ground with stones (Fig. S1). Inside each trap, we deployed three pan traps (sauce cups painted yellow, blue, and white) filled 2/3rds with soapy water (blue Dawn ® dish soap) that attract and trap bees.

We deployed traps in early morning or late evening when bees are mostly inactive, leaving them for 48 hours during fair-weather days with midday temperatures >21° C (Pane and Harmon-Threatt, 2017). In the lab, we identified bees to species (Gibbs, 2010; Ascher & Pickering, 2024). To avoid overestimating nests of social species, we excluded bees caught from the same nest during observations and only included one individual per species per treatment plot per day from emergence traps, removing 16 individuals from analyses (Table S2).

We focused on nest-making, non-parasitic, female ground-nesting bees, excluding five cleptoparasitic females and 22 males (Table S2) from analyses. Males, which do not nest, may have been captured in emergence traps waiting near female nests for mating opportunities (Visscher and Danforth, 1993). Additionally, to confirm that females were nest-making and not emerging, we noted if individuals were carrying pollen. Although, if caught using emergence traps, females may not have pollen as they have not foraged for the day. Of the ground-nesting females used for analysis, 93% had at least some pollen on them, and those that did not were exclusively from emergence traps. Thus, we can confirm that majority of females in this study were actively nesting in our plots.

### 2.5 Statistical Analyses

We tested whether environmental conditions were influenced by soil surface treatments and whether ground-nesting bee colonization (the number of nesting females) were influenced by (1) soil surface treatments and environmental soil conditions, and (2) urbanization and local site conditions.

All analyses were performed in R version 4.4.1 (R Core Team, 2024). We used package “ggplot2” (Wichham, 2016) and “ggeffects” (Lüdecke, 2018) for figures. To test variable correlationed, we used the “corrplot” package (Wei & Simko, 2024) and excluded variables with Pearson coefficients > 0.5. We assessed zero inflation and residual normally using the “Dharma” package (Hartig, 2020).

To test whether environmental conditions (soil temperature, moisture, and number of medium soil compaction readings) differed among soil surface treatments, we ran a multivariate generalized linear mixed model with “site” as a random effect with the “glmTMB” package. We then performed post-hoc univariate GLMMs for each response and Tukey pairwise comparisons using the “emmeans” package (Lenth, 2025).

We separately modeled the effects of environmental conditions (soil temperature, moisture, and number of medium soil compaction readings) and treatments on bee colonization using negative binomial generalized linear mixed models (GLMM) with “site” as a random effect using the “glmTMB” package. For treatment effects, the control was the reference level, resulting in estimate values based on the comparison between each treatment and the control. We performed post-hoc pairwise comparisons with a Tukey correction. We analyzed treatment and environmental effects separately to avoid confounding their impacts, because treatments directly altered environmental conditions.

To identify relevant predictor variables, we performed a Least Absolute Shrinkage and Selection Operator (LASSO) regression with cross-validation via the “glmnet” package (Friedman et al., 2010), which excluded moisture. The final model contained the fixed effects of temperature, number of medium compaction readings, and their interaction. For the analyses above, we aggregated data across six site visits by summing nesting females and the number of readings in each soil compaction category and averaging all other explanatory variables per subplot (15 per site). We calculated the marginal R^2^ value using the “MuMIn” package (Bartoń, 2024).

To examine treatment preferences among bee genera we used a Fisher’s Exact test with a Monte Carlo Simulation (10,000 iterations) for genera with > 10 individuals from the “stats package” (R Core Team, 2024). For genera with over 25 individuals, we conducted chi-squared goodness-of-fit tests to investigate genus-specific preferences.

We analyzed site-level conditions on bee colonization with a negative binomial generalized linear model using the “MASS” package (Venables & Ripley, 2002). We calculated McFadden’s R^2^ value using the “performance” package (Lüdecke et al., 2021). We aggregated data by site, summing nesting females and the number of readings in soil compaction categories and averaging other variables (Table 1). One site, contributing 38% of nesting bees in analyses (58/154), was an outlier (Figure S1). Thus, we log-transformed the response variable (log(y+1)) for figures for better data visualization. We performed a LASSO regression to select explanatory variables (Table 1), removing ground-nesting bee community abundance and floral abundance. We had two comparable GLM models, one that included clay and hard soil compaction, the other with sand and soft soil compaction. We chose to show the results of the clay and hard soil compaction here, based on the slightly stronger effects of these variables from the LASSO (Table S3).

**Table 1:**
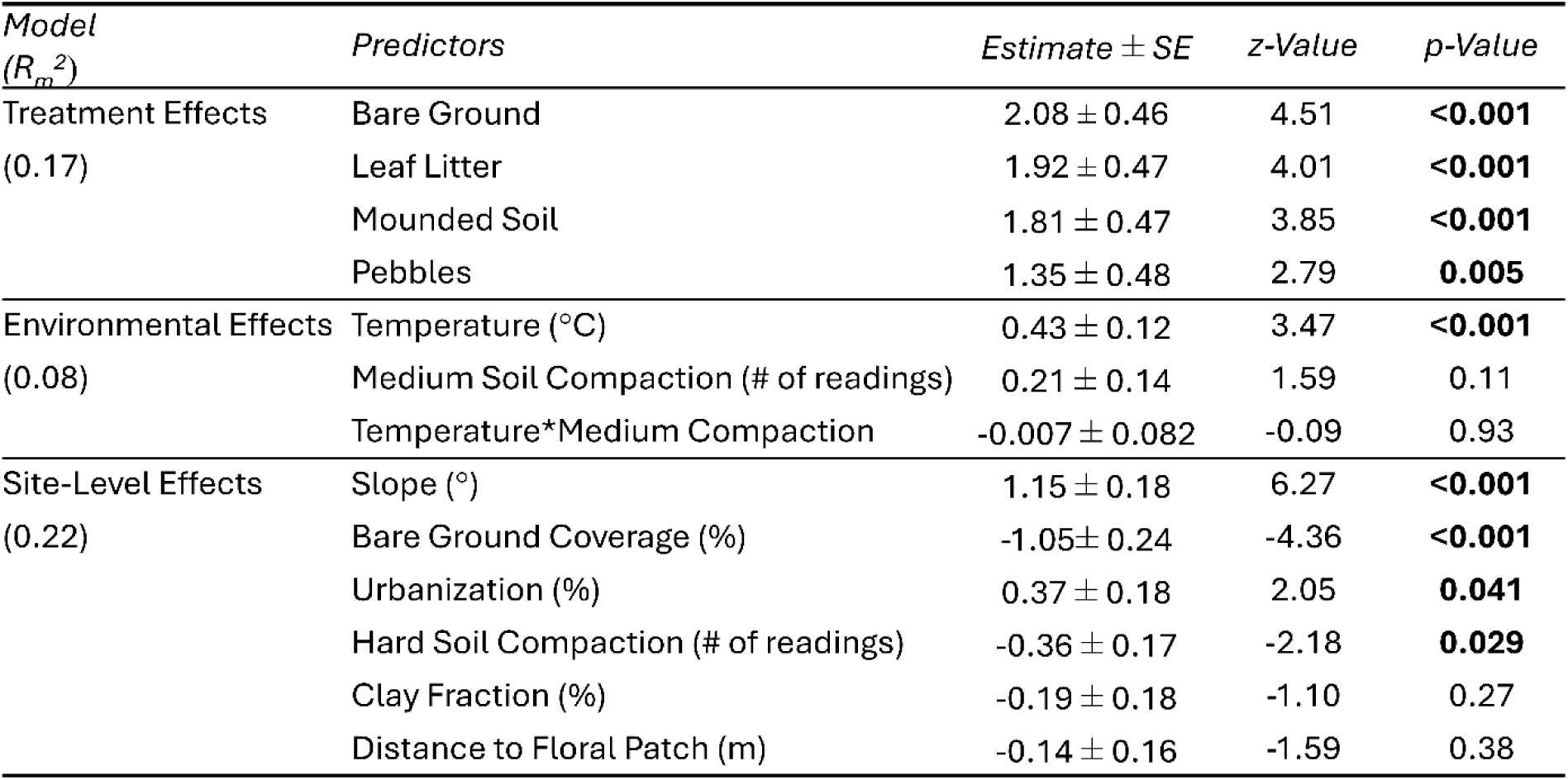
Variables with descriptions collected for site-level analysis.

## 3 Results

We collected 170 ground-nesting females from 26 species across eight genera nesting in the constructed nesting plots at 18 of 21 sites (Table S4). Eighty-one percent of ground-nesting females belonged to the family Halictidae, 16% to Andrenidae, and 2% to Apidae (Fig. S3). We also noted considerable diversity of nest entrance appearance (Fig. 2).

**Figure 2.**
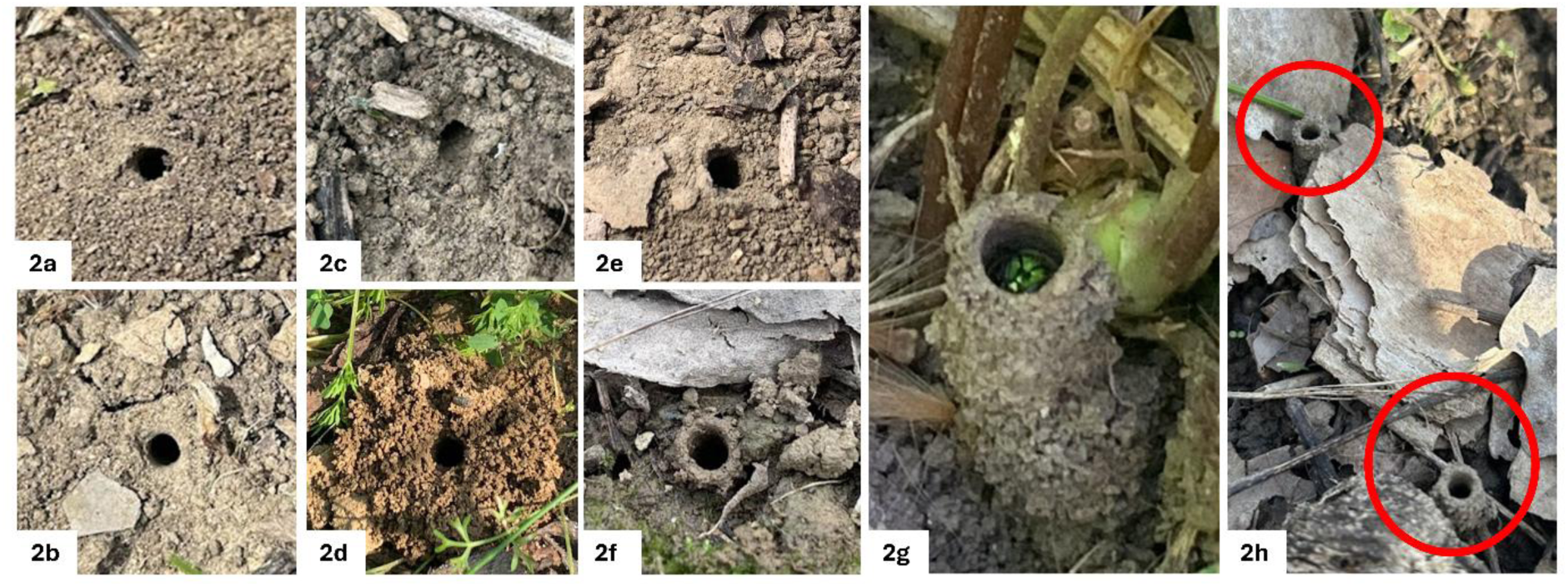
Diversity of nest hole appearance known to belong to bees during this study: 4a-4d were *Halictus* nests, 4e was a *Melissodes* nest, and 4f-4h were *Augochlorella* nests.

### 3.1 Treatment and Environmental Conditions – Hypothesis 1

We found that compared to controls, soil surface manipulations significantly increased the number of nesting bees (Table 2), and the mean number of bees found in each treatment was significantly higher than controls (Fig. 3). For environmental effects on bees, temperature was positively associated with nesting bees and there was no effect of the number of medium soil compaction readings or these variable’s interaction (Table 2). Bee abundance peaked in the medium soil compaction category (soft=30, medium=93, hard=31; Fig. S4).

**Figure 3.**
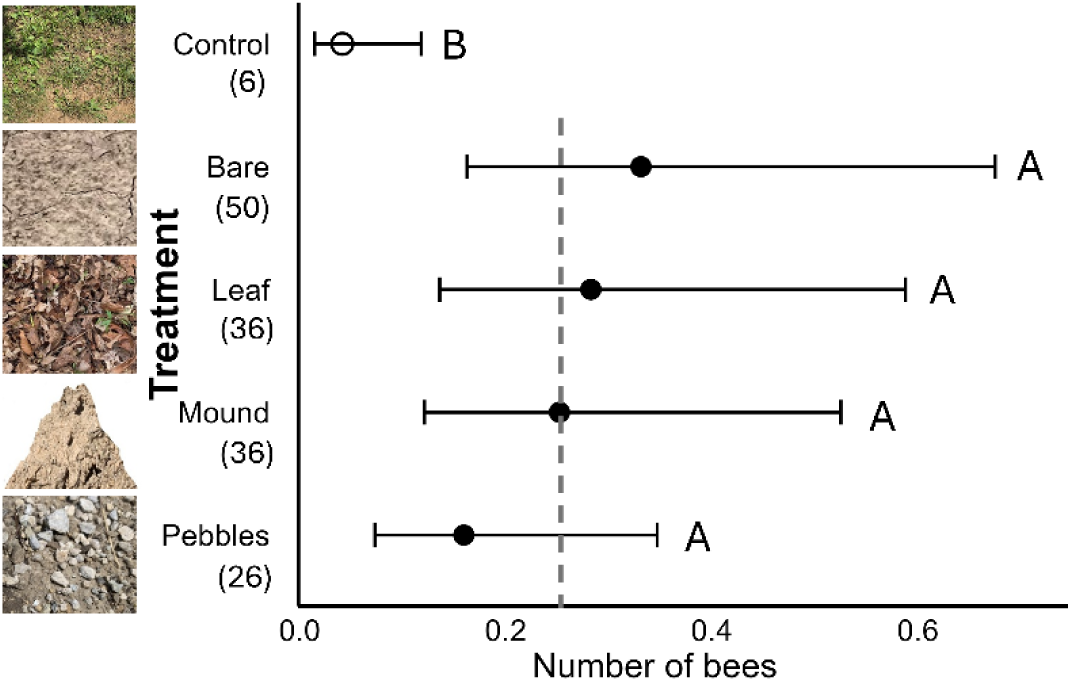
The effects of the soil surface treatments on mean bees per replicate with 95% confidence intervals. Treatments that share a letter are not significantly different based on Tukey HSD. Numbers in parentheses are total bees collected per treatment and the dotted line is the mean for all manipulated treatments (0.26).

**Table 2:**
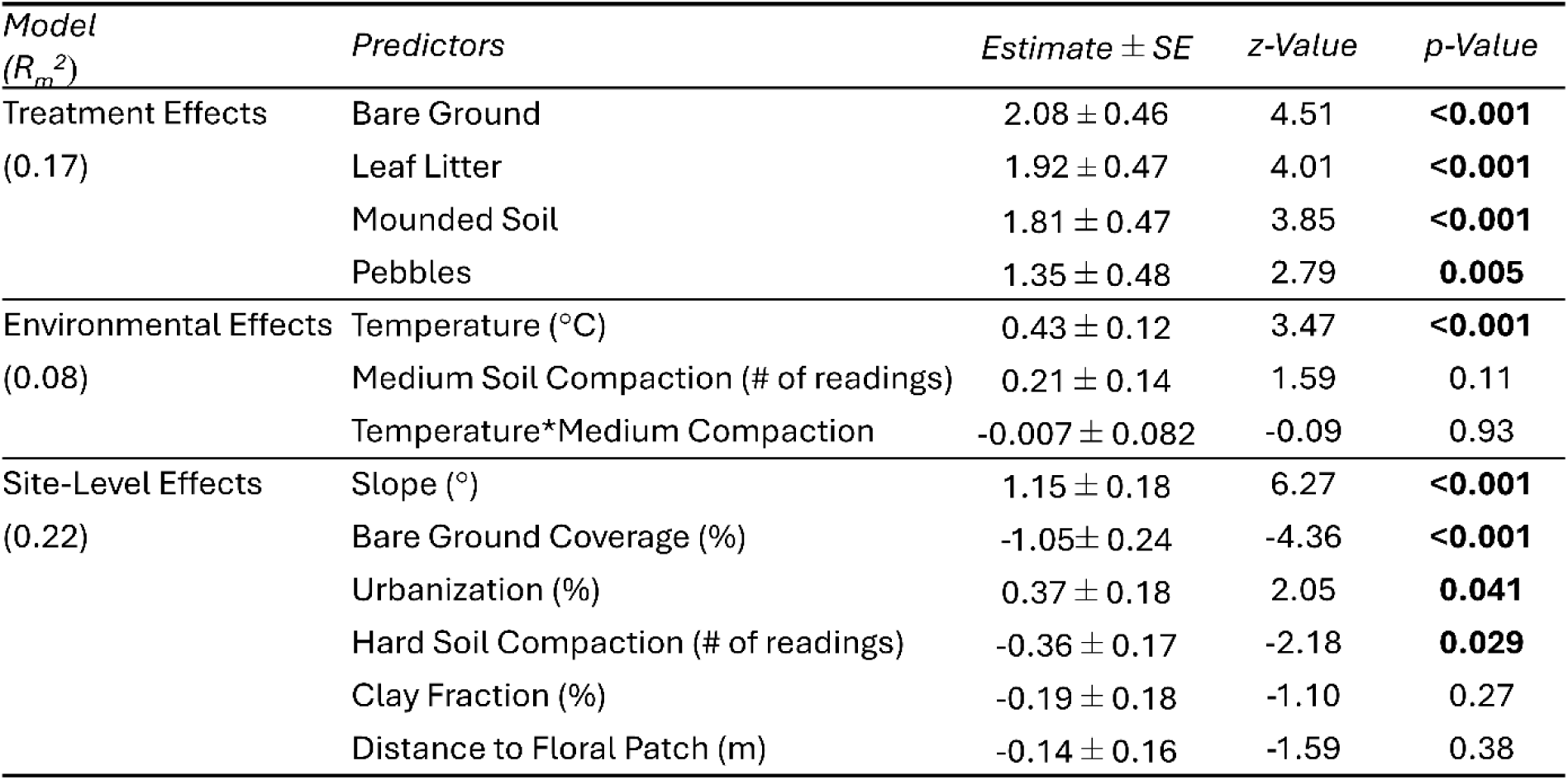
Model effects of treatments (reference level = “Control”), environmental conditions, and site conditions on bee colonization including estimate, standard error, z-value, and p-value. Marginal R_m_*^2^* presented for treatment and environmental effects and McFadden’s R*^2^* presented for site-level effects.

The multivariate mixed-effects model showed treatment type affected environmental conditions (χ² = 296.59, p < 0.001). Univariate ANOVAs revealed significant effects on temperature (F_4, 310_=60.83, p <0.001), moisture (F_4, 310_=15.24, p <0.001), and number of medium soil compaction readings (F_4, 310_=9.61, p <0.001). Pairwise comparisons (Tukey’s HSD) showed mounded soil treatments had significantly higher temperatures, followed by bare ground and pebble treatments which were significantly higher than leaf litter and control (Fig.4). Leaf litter and control treatments had significantly higher moisture than other treatment types. Leaf litter treatments had significantly more medium soil compaction readings than all treatment types except pebbles, while bare ground and control treatments had more than mounded soil (Fig. 4). Mounded soil treatments mostly had soft compaction, and bare ground and control plots had hard compaction (Table S5).

**Figure 4.**
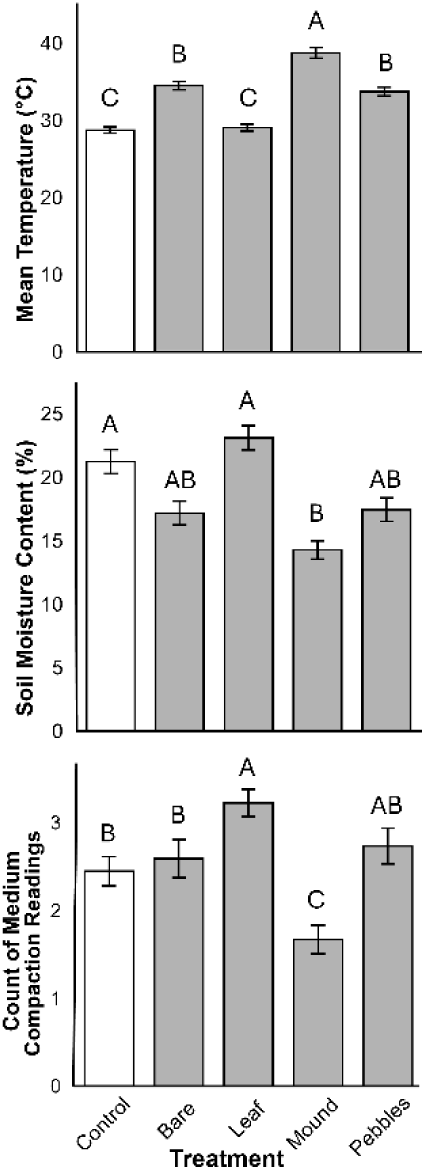
The effect of treatment per replicate on average temperature, soil moisture, and number of medium soil compaction readings (2-3.9 kg/cm^2^) with standard deviation. Treatments that share a letter are not significantly different based on Tukey HSD.

Bee genera were non-randomly associated with the soil surface manipulations (Fisher’s Exact Test: p = 0.0048). *Halictus* Latreille were found most often in the bare ground treatments (χ² = 36.9, df= 4, p<0.001). *Lasioglossum* Curtis were found relatively equally in all treatments except the control (χ²= 7.28, df=4, p=0.12). *Andrena* Fabricius was found most often in the mound and leaf litter treatments and *Augochlorella* Sandhouse in the leaf litter treatments, but these genera did not have a sample size large enough for a chi-squared test (Fig. 5).

**Figure 5.**
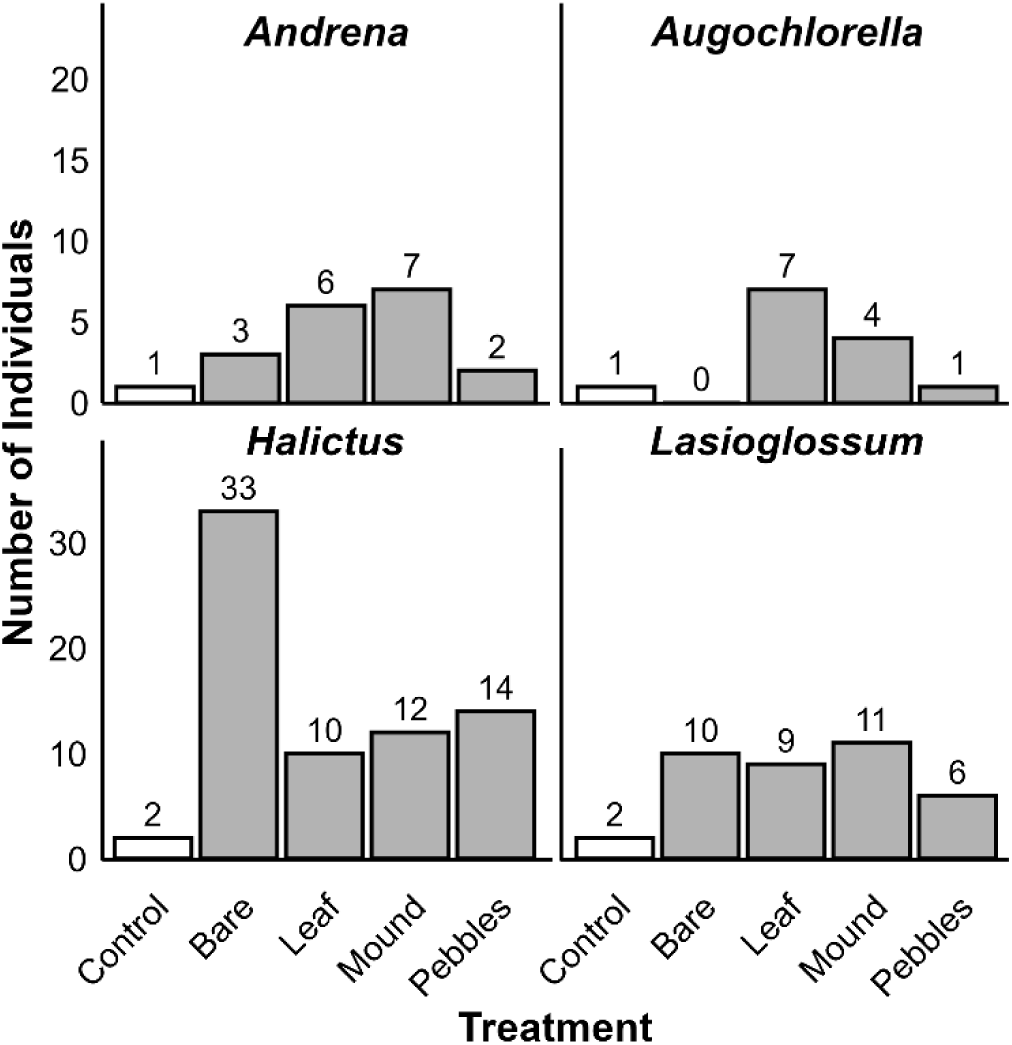
The number of individuals of the four most abundant bee genera found nesting in each soil surface treatment type.

### 3.2 Urbanization and Local Habitat Conditions – Hypothesis 2

Site-level conditions significantly affected nesting bees (McFadden’s R^2^=0.22). Sites with greater bare ground availability and number of hard soil compaction readings had fewer nesting bees. In contrast, slope and urbanization increased the number of nesting bees (Table 2: Fig. 6). Clay fraction and distance to the flower patch had no effect (Table 2).

**Figure 6:**
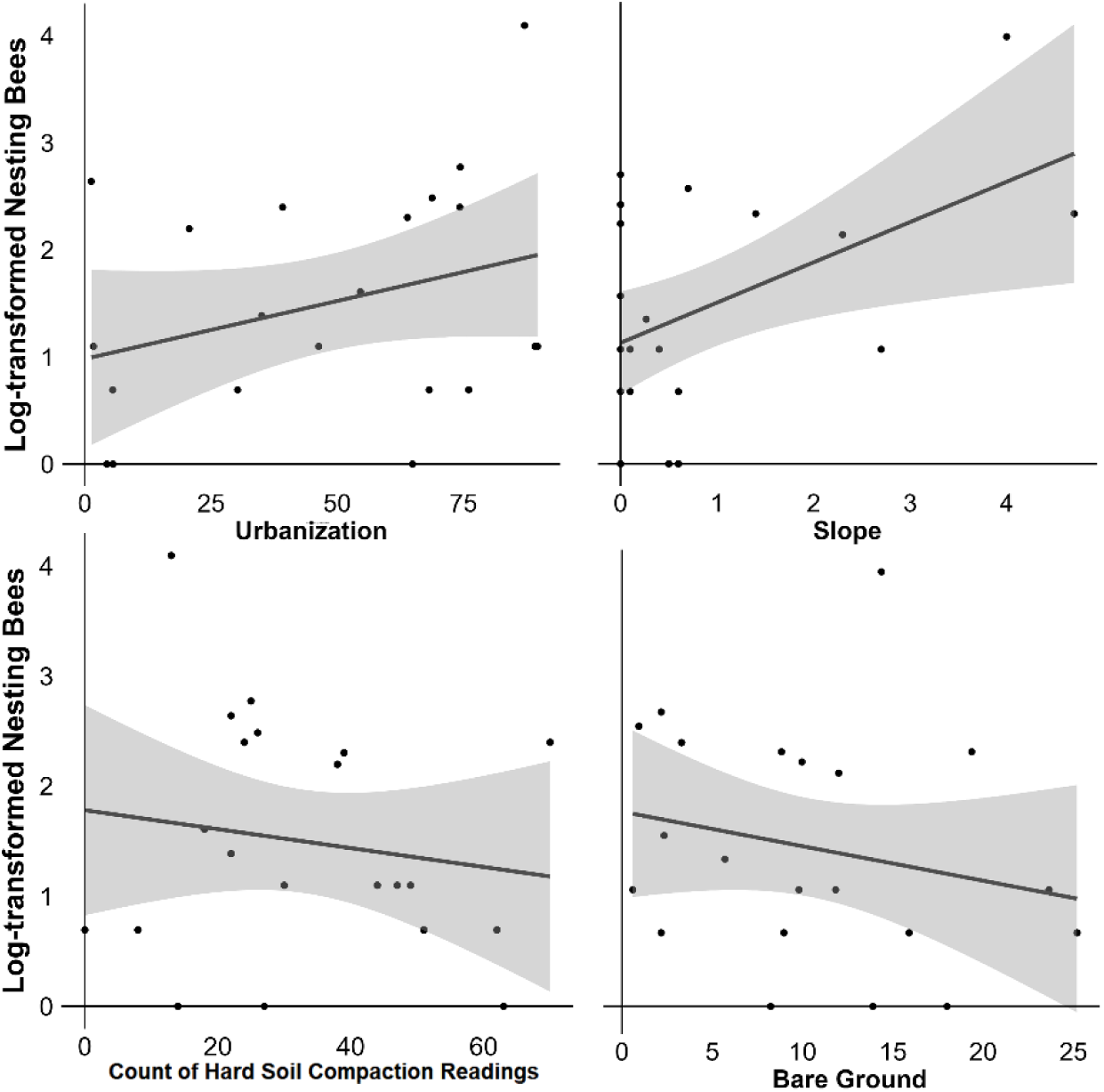
The site-level effects of urbanization, slope, number of hard soil compaction readings, and bare ground on the number of nesting bees.

## 4 Discussion

Interest in enhancing habitat for pollinator conservation is growing, but evidence-based guidance is needed. Our experimental approach showed that bees are attracted to simple soil surface manipulations. Removing vegetation and adding surface features that altered the environmental conditions revealed genus-specific substrate preferences even within bee families. These results underscore that to support diverse bee taxa, nesting habitats need to be considered at a fine taxonomic scale. Our results also point to a combination of conditions, at both site and plot level, that influence bee nesting. The high colonization of our plots at sites with little bare ground coverage and sites with high urbanization suggest that manipulations would be especially effective in sites with limited nesting habitat. Our findings provide practical guidance for attracting ground-nesting bee genera that are broadly distributed across North America and other continents and are likely generalizable to other locations.

### 4.1 Treatment and Environmental Conditions – Hypothesis 1

#### Overall patterns

Though a few studies have experimentally tested soil surface manipulations to attract ground-nesting bees (Gregory and Wright, 2005; Cane, 2015; Fortel et al, 2016; Gardein et al., 2022), ours is first to compare female nesting preferences to multiple surface soil substrates. Bees preferentially colonized all manipulated treatments over the control, indicating these substrates offered more attractive nesting sites than unaltered site conditions. Bare ground was one of multiple preferred substrates, supporting previous observations that emphasize its importance for ground-nesting bees (Potts et al., 2005; Sardiñas & Kremen, 2014). Studies documenting bare ground associations have been critiqued for potential detection bias (Harmon-Threatt, 2020), but our emergence trap data comparing nesting female abundance to the control is not subject to observer bias and validates this pattern. Notably, we show that other substrates are equally as attractive to nesting female bees as bare ground, expanding the range of effective habitat enhancements.

Bee attraction to soil surface treatments likely reflects how manipulations altered environmental conditions, but bees’ response to these conditions were complex. Nest sites with high temperatures are thought to promote early morning foraging (Stone, 1994) and have been associated with bee nesting (Potts & Willmer, 1997). However, this preference is likely species-specific and context-dependent (Maher et al., 2019). For example, mounded soil and leaf litter had similar colonization rates despite differing in temperature, possibly due to seasonal preferences. Colonization of mounded soil peaked in April (Table S6), suggesting a preference for higher temperatures during cooler months. Though warmer temperatures overall increased nesting, treatment type explained more variation, indicating that more than just temperature influences nest site preference.

Soil moisture and compaction varied by treatment and likely play a role in nest site selection, but like temperature, bee responses were not straight forward. Well-drained soils that prevent larval desiccation are expected to be preferred (Potts & Willmer, 1997), but we found no overall effect of moisture on colonization. High colonization occurred in both wetter and drier treatments. Interestingly, many individuals nested in the wetter leaf litter plots even though the nest waterproofing behaviors of many species suggest that high soil moisture is unfavorable (Cane, 1981; Visscher et al., 1994). Similarly, moderate compaction that aids in nest excavation and integrity is expected to be preferred, but optimal compaction levels remain unclear and likely differ among species (Potts & Willmer, 1997; Sardiñas & Kremen, 2014; Tsiolis et al., 2022). Colonization was three times higher in plots with medium soil compaction than other compaction categories (Fig. S4), but leaf litter plots which had more medium compaction readings were not colonized more than other manipulated treatments. These results suggest bees can tolerate a range of moisture and compaction and highlight that nest site selection is shaped by multiple factors rather than any single environmental condition.

A commonality between all manipulated treatments was a lack vegetation which likely made them more attractive for nesting, while control plots with intact vegetation had very low colonization. This supports prior findings that dense vegetation can inhibit nesting by preventing excavation (Packer & Knerer, 1986; Wuellner, 1999). Although nests are harder to locate in vegetation (Harmon-Threatt, 2020), our use of emergence traps confirms that bees actively avoided vegetated plots. The absence of bees found in controls also suggests the high colonization in other treatments resulted from attraction to plots, not preexisting nests. Tilling to remove vegetation may have altered soil compaction initially, but treatments differed in compaction. This suggests that tilling was not the primary factor driving colonization and further supports the influence of vegetation. Whether bees are drawn to disturbed soil remains an open question that should be investigated in the future.

Beyond environmental conditions, unmeasured factors like visual cues and parasite protection might have influenced nest site selection. Features that help bees relocate nest entrances or conceal them from cleptoparasites provide advantages (Brünnert et al., 1994; Wcislo, 1996). Mounded soil may be easier to spot from a distance and bare ground offers exposed entrances for short-range visibility. Pebbles can offer both visual cues and concealment, but surprisingly, these treatments did not attract more bees than bare ground, in contrast to results from Cane (2015) where pebbles doubled nest numbers. While some bees may prefer increased visibility, leaf litter may be attractive because it helps conceal nest entrances from cleptoparasites that enter the unguarded nests while the female forages (Visscher and Danforth, 1993). However, the strong association with bare ground suggests that not all species respond to this risk in the same way.

#### Genus-Specific Patterns

The most abundant bee genera in this study preferred different soil surface treatments. *Halictus* and *Lasioglossum* were most frequently captured, consistent with studies showing vegetation removal attracts these species (Gregory & Wright, 2005; Fortel et al., 2016; Tsiolis et al. 2022). These primitively social species produce multiple generations annually and can reach high abundances (Michener, 2007). In our study, *Halictus* preferred bare ground treatments, while *Lasioglossum* showed no clear treatment preference. *Lasioglossum* is more diverse compared to *Halictus* and the preferences seen here likely reflect the highly abundant species *Halictus ligatus* Say, 1837 (76% of *Halictus* individuals). Thus, nesting preferences may be specific-specific rather than genera-specific.

Less abundant genera also showed substrate preferences. *Andrena* primarily occupied leaf litter and mounded soil treatments, while *Augochlorella* primarily occupied leaf litter treatments. Most *Andrena* species are active in the spring, forage in forests, and are subject to high levels of parasitism (Alexander et al., 1991). Abundant leaf litter in forests offer protection against cleptoparasites, and mounded soil offers warmer environmental conditions in the cooler spring months. *Augochlorella* preferred leaf litter treatments, where they often built visible turrets (Fig. 4h), perhaps to aid nest relocation while nesting in preferrable environmental conditions. Our small sample size of *Andrena* and *Augochlorella* preclude further analyses of but highlight patterns that merit further research. Interestingly, the genera *Halitus*, *Lasioglossum*, and *Augochlorella* all belong to the family Halictidae, highlighting that genera in the same family have distinct nesting preferences.

### 4.2 Urbanization and Local Habitat Conditions – Hypothesis 2

We show that limited nesting habitat at both landscape and local scales influenced colonization. Although urbanization reduces ground-nesting bee richness and abundance, higher proportions of impervious surface mean less area of suitable nesting habitat (Hernandez et al., 2009; Ayers & Rehan, 2021). As predicted, colonization was higher in urban sites. Colonization was also higher at sites with less bare ground, supporting the influence of local site conditions on pollinator habitat use (Burks & Philpott, 2017). These results suggest that in highly urban and vegetated sites with limited bare ground constructed nest plots would be a valuable tool for conservation.

Additional site-level factors, including slope and hard soil compaction affected bee colonization. Sites with predominantly hard-compacted soils supported fewer nests, suggesting that high soil compaction may act as a barrier to nesting in these habitat enhancements. This finding indicates that soil surface manipulations may only be effective when implemented in areas with softer soil compaction or with a variety of compaction levels which can provide more suitable nesting opportunities. Additionally, while we placed plots on relatively flat terrain (variance of 0 to 4 degrees), even this small range influenced nest site colonization. Sloped ground may offer benefits such as increased sun exposure and water drainage. Many species nest in vertical or steep slopes (Norden, 1984; Sardiñas & Kremen, 2014), and our findings show that even shallow slopes can enhance nest site suitability for some species.

We detected no effect of distance to the flower patch or soil texture. Distance to flower patch is expected to affect bees because closer nests improve foraging efficiency (Zurbachen et al, 2010). However, we located all experimental plots within 200 m of a flower patch and this short distance may not reduce foraging efficiency. Similarly clay fraction had no effect, consistent with other studies (Fortel et al., 2016). Previous studies suggest bees may prefer sandy soils (Cane, 1991; López-Uribe et al., 2015). It is important to note that sand had no effect in the alternate model (Table S3). However, sand may still be important for species that specialize in sandy soils which were absent in our study, or in areas with a wider range of sand content.

## 5 Conclusions

This study demonstrates that simple soil surface manipulations, or soil sanctuaries, such as constructing bare ground patches, adding pebbles, forming soil mounds, and incorporating leaf litter near floral resources can enhance nesting habitat for ground-nesting bees. Our experimental approach along with using both emergence traps and observations validates and builds on patterns seen in observational studies. We show that bees preferred manipulated treatments over controls, with treatment preferences varying by genus, and that nest site choice is multi-faceted. Environmental conditions at the plot-level such as higher temperature increased nesting activity while site-level conditions such as sloped ground and urbanization also increased colonization, and hard soil compaction and bare ground availability reduced it. These findings highlight the importance of tailoring nesting enhancements to local site conditions– e.g., loosening compacted soil or constructing habitats on slopes improve habitat suitability. The increased nesting at urban sites and sites with less bare ground suggest that these soil sanctuaries effectively provide nesting sites where natural options are limited, supporting the need to enhance nesting habitat across a variety of landscapes.

Our results contribute to the growing evidence that diverse substrates can support ground-nesting bees in both natural and human-altered landscapes. As pollinator gardens become more common, integrating suitable nesting habitat is essential. Soil sanctuaries offer a practical strategy for land managers and gardeners alongside pollinator gardens. In disturbed environments such as urban or agricultural areas, and in natural environments where nesting may be limited due to high vegetation density, soil sanctuaries can create stable, protected nesting sites. Maintaining these habitats for multiple years may yield long-term demographic benefits by fostering local population growth (Roulston & Goodell, 2011). This research illustrates how small-scale habitat enhancement can have meaningful ecological benefits by supporting biodiversity. It also emphasizes the need for ecological refugees that promote pollinator persistence and population resilience amid ongoing environmental change.

## Acknowledgements

We would like to thank Rex Harvey-Thurston, Abby Subler, and Lizzy Sakulich for field assistance, MaLisa Spring for assistance identifying bee specimens, Joey Smith for assistance on soil texture analysis, Laura Fey for connecting us with land managers, and all who provided feedback on this manuscript. We thank all the cooperating land management agencies for allowing us to conduct our experiment. This work was funded by The Garden Club of America Board of Associates Centennial Pollinator Fellowship.

## Author Contributions

A. F. and K. G. conceived the ideas and designed methodology. A. F. collected and analyzed the data and led the writing of the manuscript. All authors contributed critically to the drafts and gave final approval for publication.

## Data Availability Statement

Data available via figshare: https://doi.org/10.6084/m9.figshare.28590200 (Fredenburg & Goodell, 2025)

## Conflict of Interest

The authors have no conflicts of interest to declare.

## Notes

https://doi.org/10.6084/m9.figshare.28590200

### Competing Interest Statement

The authors have declared no competing interest.

