## Supplementary figures and images for "Soil sanctuaries: Experimental manipulations enhance ground-nesting bee habitat across an urban gradient"

### Figure S1

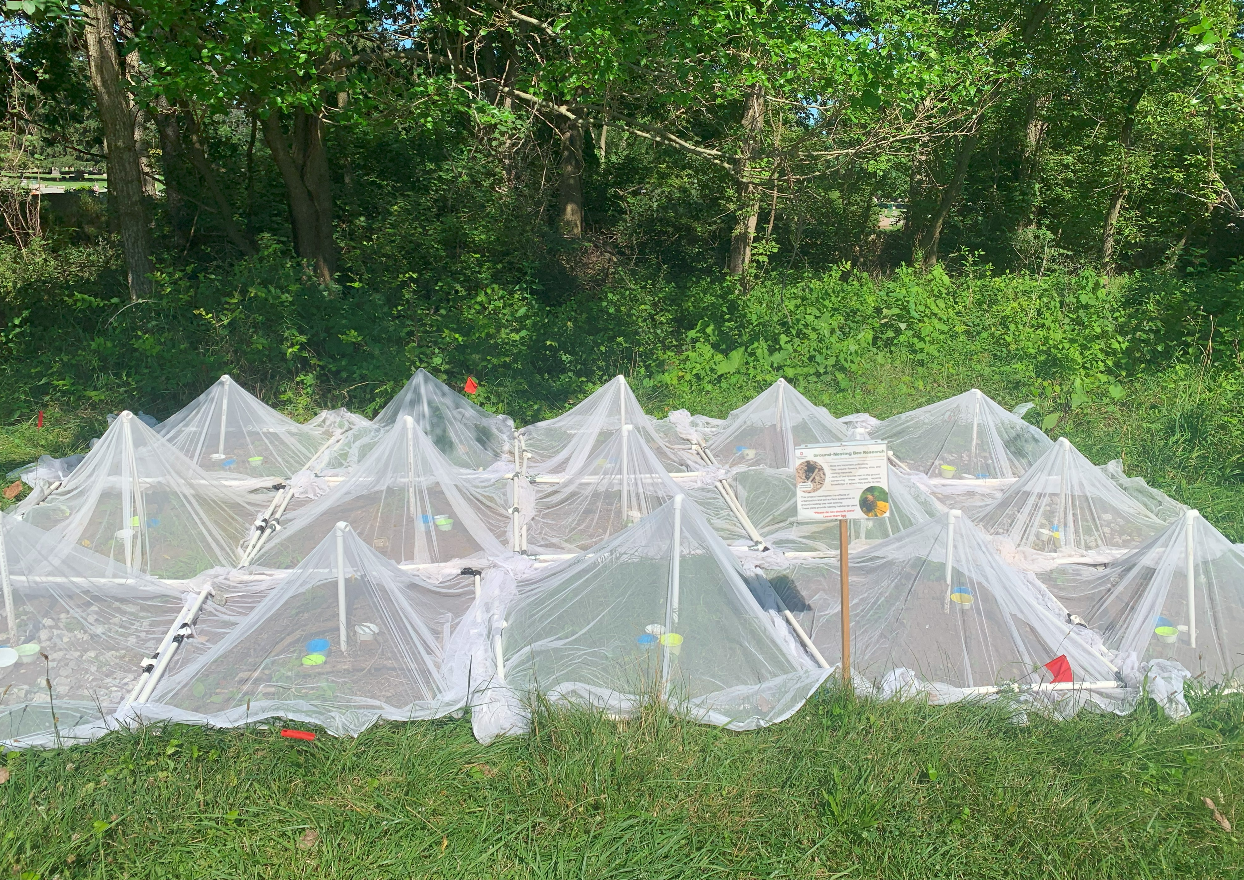

**Figure S1:** Emergence traps placed over an experimental plot.

### Figure S2

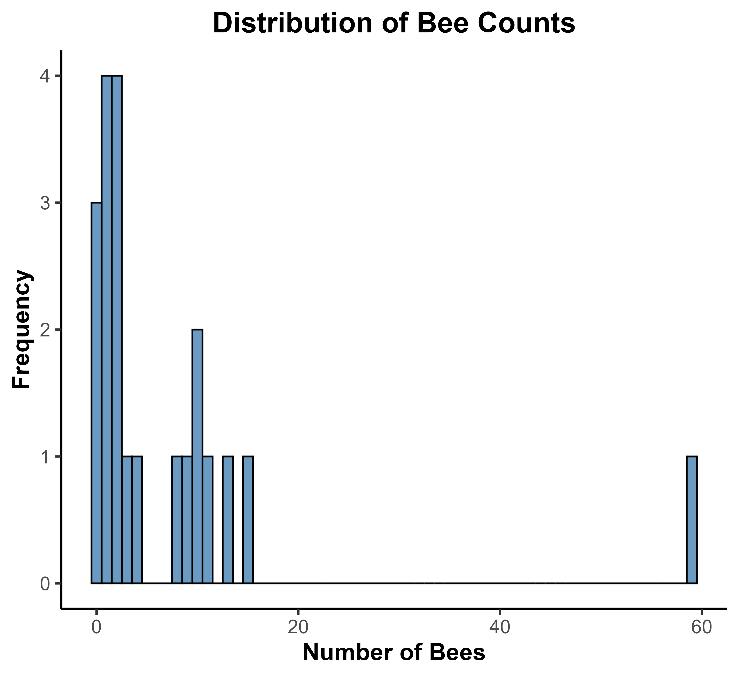

**Figure S2:** Distribution of the response variable, nesting bees, aggregated by site.

### Figure S3

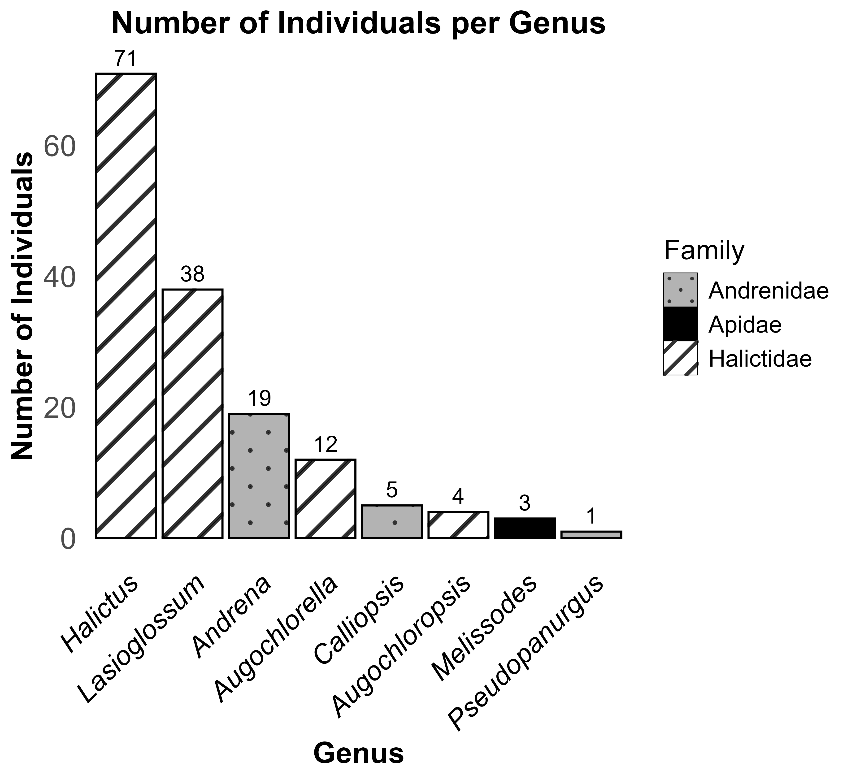

**Figure S3:** The number of female ground-nesting bees per genus caught and included in analysis.
