## Supplementary material for "Soil sanctuaries: Experimental manipulations enhance ground-nesting bee habitat across an urban gradient": Figure S4

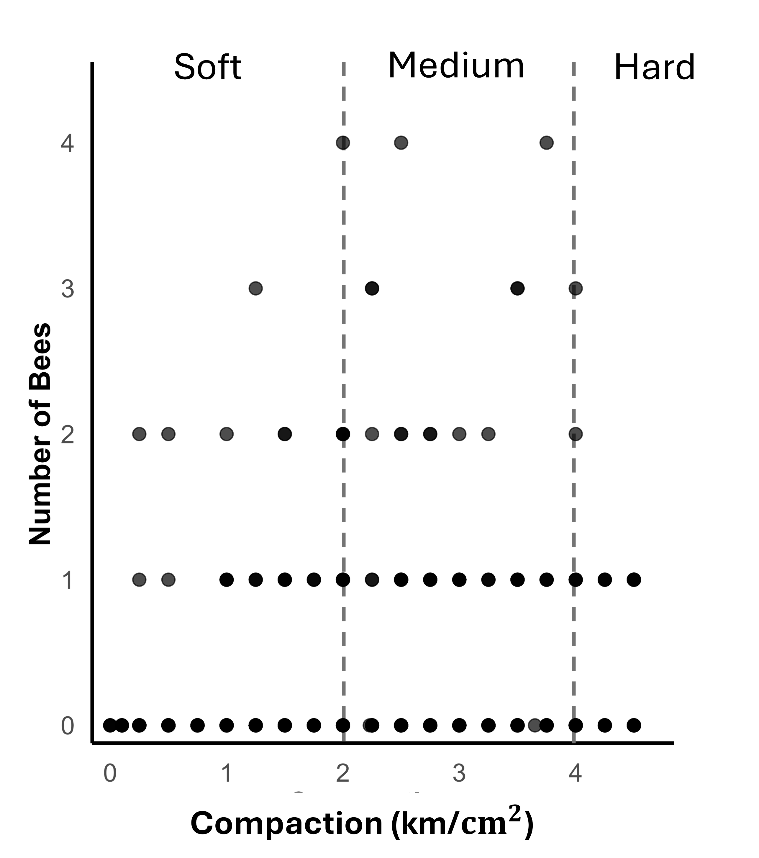

**Figure S4:** The effect of compaction as a continuous variable on nesting bees, with lines representing the categories used in analyses. Each datapoint is a compaction reading of one of the 15 1-m x 1-m treatments replicates per site at a visit.
