## Supplementary material for "Soil sanctuaries: Experimental manipulations enhance ground-nesting bee habitat across an urban gradient": Table S1

| Site Name | Latitude | Longitude |
| --- | --- | --- |
| Slate Run | 39.76178 | -82.8514 |
| Glacier Ridge | 40.15687 | -83.1959 |
| Prairie Oaks | 39.99554 | -83.2679 |
| Hoover Prairie | 40.11081 | -82.8726 |
| Chestnut Ridge | 39.80773 | -82.7558 |
| Delco Water Headquarters | 40.2026 | -83.0564 |
| Godown Park | 40.08322 | -83.0513 |
| Civic Park | 39.97031 | -82.8181 |
| Inniswood | 40.10358 | -82.8988 |
| OSU Wetland | 40.0198 | -83.02 |
| Whitney Playground | 40.10508 | -83.034 |
| Ohio School for the Deaf | 40.06954 | -83.0013 |
| OSU Olentangy | 40.00937 | -83.0194 |
| Burbank Park | 40.05321 | -83.0851 |
| Thompson Park | 40.04373 | -83.0704 |
| Woodward Nature Preserve | 40.07221 | -82.9874 |
| Clinton-Como Park | 40.02636 | -83.0224 |
| Sawmill Wetlands | 40.09882 | -83.0828 |
| Chadwick | 40.01144 | -83.0305 |
| Riverside | 40.02845 | -83.0374 |
| Scioto Audubon | 39.94575 | -83.0073 |

**Table S1:** Site Names and GPS Coordinates of each location.
