## Supplementary material for "Soil sanctuaries: Experimental manipulations enhance ground-nesting bee habitat across an urban gradient": Table S2

| **Date** | **Site Name** | **Visit** | **Replicate** | **Plot Type** | **Sex** | **Species** | **Reason for Removal** |
| --- | --- | --- | --- | --- | --- | --- | --- |
| 5/22/2023 | Clinton-Como Park | 1 | 2 | Bare | F | *Nomada sp* | Parasitic |
| 6/3/2023 | Sawmill Wetlands | 1 | 1 | Bare | F | *Halictus ligatus* | Social |
| 6/7/2023 | Chestnut Ridge | 1 | 2 | Leaf | M | *Calliopsis andreniformis* | Male |
| 6/7/2023 | Chestnut Ridge | 1 | 2 | Pebbles | M | *Calliopsis andreniformis* | Male |
| 7/14/2023 | Ohio School for the Deaf | 3 | 2 | Bare | M | *Calliopsis andreniformis* | Male |
| 7/14/2023 | Riverside | 3 | 1 | Pebbles | M | *Calliopsis andreniformis* | Male |
| 7/14/2023 | Scioto Audubon | 3 | 1 | Leaf | F | *Lasioglossum ephialtum* | Social |
| 7/17/2023 | Godown | 3 | 2 | Bare | M | *Calliopsis andreniformis* | Male |
| 7/17/2023 | Sawmill Wetlands | 3 | 3 | Bare | M | *Halictus confusus* | Male |
| 7/17/2023 | Sawmill Wetlands | 3 | 2 | Pebbles | F | *Lasioglossum versatum* | Social |
| 7/17/2023 | Sawmill Wetlands | 3 | 3 | Bare | F | *Halictus ligatus* | Social |
| 7/17/2023 | Sawmill Wetlands | 3 | 3 | Bare | F | *Halictus confusus* | Social |
| 7/19/2023 | Thompson Park | 3 | 2 | Leaf | M | *Calliopsis andreniformis* | Male |
| 7/19/2023 | Thompson Park | 3 | 3 | Pebbles | M | *Halictus rubicundus* | Male |
| 7/19/2023 | Thompson Park | 3 | 3 | Bare | M | *Calliopsis andreniformis* | Male |
| 7/20/2023 | Slate Run | 3 | 2 | Bare | M | *Calliopsis andreniformis* | Male |
| 7/20/2023 | Slate Run | 3 | 2 | Leaf | M | *Calliopsis andreniformis* | Male |
| 7/21/2023 | Delco Water | 3 | 1 | Pebbles | M | *Calliopsis andreniformis* | Male |
| 7/21/2023 | Delco Water | 3 | 2 | Control | F | *Halictus ligatus* | Social |
| 7/26/2023 | Slate Run | 3 | 2 | Bare | M | *Calliopsis andreniformis* | Male |
| 7/26/2023 | Slate Run | 3 | 2 | Bare | F | *Halictus ligatus* | Social |
| 8/8/2023 | Sawmill Wetlands | 4 | 3 | Bare | F | *Holcopasites calliopsidis* | Parasitic |
| 8/11/2023 | Delco Water | 4 | 1 | Leaf | M | *Calliopsis andreniformis* | Male |
| 8/11/2023 | Delco Water | 4 | 2 | Control | F | *Halictus ligatus* | Social |
| 8/11/2023 | Delco Water | 4 | 2 | Control | F | *Halictus ligatus* | Social |
| 4/15/2024 | OSU Wetlands | 6 | 2 | Bare | F | *Nomada Bidentate sp* | Male |
| 4/18/2024 | Whitney Playground | 6 | 1 | Mound | M | *Andrena nasonii* | Male |
| 4/18/2024 | Whitney Playground | 6 | 1 | Leaf | F | *Augochlorella aurata* | Social |
| 4/18/2024 | Whitney Playground | 6 | 1 | Leaf | F | *Augochlorella aurata* | Social |
| 4/18/2024 | Sawmill Wetlands | 6 | 3 | Pebbles | F | *Lasioglossum versatum* | Social |
| 4/18/2024 | Sawmill Wetlands | 6 | 1 | Pebbles | F | *Halictus ligatus* | Social |
| 4/18/2024 | Sawmill Wetlands | 6 | 2 | Mound | F | *Halictus confusus* | Social |
| 4/18/2024 | Sawmill Wetlands | 6 | 2 | Bare | F | *Halictus confusus* | Social |
| 4/18/2024 | Sawmill Wetlands | 6 | 2 | Bare | F | *Lasioglossum versatum* | Social |
| 4/29/2024 | Ohio School for the Deaf | 6 | 1 | Pebbles | M | *Andrena cressonii* | Male |
| 4/29/2024 | Ohio School for the Deaf | 6 | 2 | Mound | M | *Andrena miserabilis* | Male |
| 4/29/2024 | Ohio School for the Deaf | 6 | 2 | Pebbles | M | *Andrena cressonii cressonii* | Male |
| 4/29/2024 | Ohio School for the Deaf | 6 | 2 | Leaf | M | *Andrena cressonii cressonii* | Male |
| 4/29/2024 | Ohio School for the Deaf | 6 | 2 | Leaf | M | *Andrena cressonii cressonii* | Male |
| 4/29/2024 | Godown | 6 | 3 | Mound | M | *Andrena nasonii* | Male |
| 5/1/2024 | Glacier Ridge | 6 | 1 | Leaf | M | *Andrena gardinari* | Male |
| 5/8/2024 | Hoover Prairie | 6 | 1 | Leaf | F | *Nomada Bidentate sp* | Parasitic |
| 5/8/2024 | Hoover Prairie | 6 | 1 | Leaf | F | *Nomada Bidentate sp* | Parasitic |

**Table S2:** List of all bee individuals omitted from the analyses with justification for omission.
