## Supplementary material for "Soil sanctuaries: Experimental manipulations enhance ground-nesting bee habitat across an urban gradient": Table S3

| *Model*  *(R_m_^2^*) | *Predictors* | *Estimate ± SE* | *z-Value* | *p-Value* |
| --- | --- | --- | --- | --- |
| Site-Level Effects | Slope (°) | 1.16 *±* 0.17 | 6.94 | **<0.001** |
| (0.23) | Bare Ground Coverage (%) | -1.16 *±* 0.22 | -5.30 | **<0.001** |
|  | Urbanization (%) | 0.42 *±* 0.16 | 2.70 | **0.007** |
|  | Soft Soil Compaction (# of readings) | 0.43 *±* 0.18 | -1.64 | **0.020** |
|  | Sand Fraction (%) | 0.24 *±* 0.14 | 1.64 | 0.10 |
|  | Distance to Floral Patch (m) | -0.22 *±* 0.14 | -1.59 | 0.11 |

**Table S3:** Alternative site-level model including sand fraction and soft soil compaction.
