## Supplementary material for "Soil sanctuaries: Experimental manipulations enhance ground-nesting bee habitat across an urban gradient": Table S4

| **Species Name** | **Bare** | **Control** | **Leaf** | **Mound** | **Pebbles** | **Species Total** |
| --- | --- | --- | --- | --- | --- | --- |
| Andrena algida | 1 |  |  |  |  | 1 |
| Andrena cressonii cressonii | 1 |  |  |  |  | 1 |
| Andrena erigeniae | 1 |  | 2 | 3 |  | 6 |
| Andrena ilicis |  | 1 |  | 1 |  | 2 |
| Andrena imitatrix |  |  | 1 | 2 | 1 | 4 |
| Andrena robertsonii |  |  | 1 | 1 |  | 2 |
| Andrena violae |  |  | 2 |  | 1 | 3 |
| Augochlorella aurata |  | 1 | 7 | 3 | 1 | 12 |
| Augochlorella sp |  |  |  | 1 |  | 1 |
| Augochloropsis metallica |  |  |  | 1 |  | 1 |
| Augochloropsis viridula | 1 |  | 2 |  |  | 3 |
| Calliopsis andreniformis | 2 |  | 1 | 1 | 1 | 5 |
| Halictus confusus | 5 |  | 2 | 3 | 2 | 12 |
| Halictus ligatus | 24 | 2 | 8 | 9 | 11 | 54 |
| Halictus rubicundus | 4 |  |  |  | 1 | 5 |
| Lasioglossum anomalum | 1 |  |  | 1 |  | 2 |
| Lasioglossum ephialtum |  |  | 1 | 1 | 1 | 3 |
| Lasioglossum hitchensi | 1 | 1 | 5 | 1 | 3 | 11 |
| Lasioglossum illinoense | 1 | 1 |  | 1 |  | 3 |
| Lasioglossum imitatum |  |  | 1 |  |  | 1 |
| Lasioglossum paradmirandum |  |  | 1 | 1 |  | 2 |
| Lasioglossum pilosum | 1 |  |  | 3 |  | 4 |
| Lasioglossum sp | 3 |  | 1 | 1 |  | 5 |
| Lasioglossum tegulare | 1 |  |  | 1 | 1 | 3 |
| Lasioglossum versatum | 2 |  |  | 1 | 1 | 4 |
| Melissodes subillatus |  |  |  |  | 2 | 2 |
| Melissodes trinodis | 1 |  |  |  |  | 1 |
| Pseudopanurgus labrosiformis |  |  | 1 |  |  | 1 |
| **Plot Type Total** | **50** | **6** | **36** | **36** | **26** | **154** |

**Table S4:** List of all bee species and their abundances found in each treatment type and in total.
