## Supplementary material for "Soil sanctuaries: Experimental manipulations enhance ground-nesting bee habitat across an urban gradient": Table S5

| *Variables* | *Bare* | *Control* | *Leaf* | *Mound* | *Pebbles* |
| --- | --- | --- | --- | --- | --- |
| Temperature | 34.46 ± 0.54 | 28.7 ± 0.49 | 29.02 ± 0.44 | 38.71 ± 0.68 | 33.69 ± 0.57 |
| Moisture | 17.16 ± 0.94 | 21.21 ± 0.009 | 23.09 ± 0.96 | 14.25 ± 0.72 | 17.43 ± 0.92 |
| Medium Compaction | 2.59 ± 0.22 | 2.44 ± 0.17 | 3.22 ± 0.15 | 1.67 ± 0.16 | 2.73 ± 0.20 |
| Soft Compaction* | 0.43 ± 0.13 | 0.22 ± 0.10 | 1.29 ± 0.16 | 3.79 ± 0.21 | 0.63 ± 0.12 |
| Hard Compaction* | 2.98 ± 0.24 | 3.33 ± 0.20 | 1.49 ± 0.14 | 0.54 ± 0.16 | 2.63 ± 0.24 |
| Continuous Compaction (kg/cm^2^)* | 3.63 ± 0.09 | 3.82 ± 0.08 | 2.88 ± 0.08 | 1.71 ± 0.12 | 3.48 ± 0.10 |

**Table S5:** Mean and standard error of environmental conditions from each treatment type from a replicate including number of medium, soft, and hard soil compaction readings, compaction (kg/cm^2^), temperature (°C), and moisture (%). *Other compaction metrics were not used in analyses but provided here for result interpretation.
