## Supplementary material for "Soil sanctuaries: Experimental manipulations enhance ground-nesting bee habitat across an urban gradient": Table S6

| ***Month*** | ***Bare*** | ***Control*** | ***Leaf*** | ***Mound*** | ***Pebbles*** | ***Month Total*** |
| --- | --- | --- | --- | --- | --- | --- |
| April | 13 | 3 | 9 | 20 | 10 | **55** |
| May | 6 | 0 | 3 | 3 | 3 | **15** |
| June | 7 | 1 | 1 | 2 | 4 | **15** |
| July | 13 | 1 | 12 | 6 | 6 | **38** |
| August | 10 | 1 | 10 | 5 | 5 | **29** |
| September | 1 | 0 | 1 | 0 | 0 | **2** |
| ***Plot Total*** | **50** | **6** | **36** | **36** | **26** | **154** |

**Table S6:** Number of bees caught in each plot type broken down by month.
